# Staphylococcal superantigen-like protein 13 activates neutrophils via Formyl Peptide Receptor 2

**DOI:** 10.1101/305847

**Authors:** Yuxi Zhao, Kok P. M. van Kessel, Carla J. C. de Haas, Malbert R. C. Rogers, Jos A. G. van Strijp, Pieter-Jan A. Haas

## Abstract

Staphylococcal Superantigen-Like (SSL) proteins, one of major virulence factor families produced by *Staphylococcus aureus*, were previously demonstrated to be immune evasion molecules that interfere with a variety of innate immune defenses. However, in contrast to these characterized SSLs, that inhibit immune functions, we show that SSL13 is a strong activator of neutrophils via the formyl-peptide receptor 2 (FPR2). Moreover, our data show that SSL13 acts as a chemoattractant, induces degranulation and oxidative burst in neutrophils. As with many other staphylococcal immune evasion proteins, SSL13 shows a high degree of human specificity. SSL13 is not able to efficiently activate mouse neutrophils, hampering *in vivo* experiments.

In conclusion, SSL13 is a neutrophil chemoattractant and activator that acts via the FPR2. Therefore, SSL13 is a unique SSL member that does not belong to the immune evasion class, but is a pathogen alarming molecule.

## Importance

This study describes the target receptor and mechanism of action of Staphylococcal superantigen like 13 (SSL13), a secreted protein residing on *S. aureus* immune evasion cluster 2(IEC-2). In sharp contrast to other previously characterized SSLs located on the staphylococcal pathogenicity island 2 (SPI-2), that inhibit immune functions, we demonstrate that SSL13 is a chemoattractant and a neutrophil activator that acts via the FPR2. Therefore, SSL13 is a unique SSL member not belonging to the immune evasion class, but is a pathogen alarming molecule sensed by the FPR2. Our study provides a new concept of SSLs; SSLs not only inhibit host immune processes but also recruit human neutrophils to the site of infection. This new insight allows us to better understand complex interactions between host and *S. aureus* pathological processes.

## Introduction

The Gram-positive bacterium *Staphylococcus aureus* (*S. aureus*) is an opportunistic human pathogen that causes a wide range of diseases from mild skin infections to more serious life-threatening wound and systemic infections (1). In order to successfully invade and colonize the human host, *S. aureus* secretes a large arsenal of immune evasion molecules that specifically target components of the human innate and adaptive immune systems (2). These secreted proteins interfere with a range of immune defenses, which can be grouped into four categories: blocking, degradation, cell lysis and modulation (3). Despite the functional differences and diversity in targets, the staphylococcal immune evasion proteins are secreted proteins that show remarkable resemblances. These proteins contain very conserved structural properties (4). They are often small, varying in size between 8 and 35-kDa and have extreme isoelectric points (above 9 or below 5). Another common property of these proteins is that they are located on genomic clusters with other virulence factors. The secretome of *S. aureus* is predicted to contain up to 270 proteins, of which over 35 staphylococcal evasion molecules have been described (3). Identification and characterization of these secreted proteins will lead to a better understanding of the *S. aureus* pathological processes.

Neutrophils play a crucial role in protecting the host from *S. aureus* infections (5). Inherited or acquired neutrophil dysfunction, such as leukocyte adhesion deficiency and chronic granulomatous disease, lead to an increased risk of severe *S. aureus* infections (6). Disruption of physical barriers and invasion of *S. aureus* initiates the release of pro-inflammatory signals that promote neutrophil adherence to the vascular endothelium, extravasation and migration from the bloodstream towards to the site of infection (1). However, *S. aureus* can subvert neutrophil functions via the secretion of proteins that inhibit neutrophil recruitment and activation(7, 8). A variety of immune evasion proteins have been identified that specifically target neutrophil surface receptors. Some immune evasion proteins inhibit pro-inflammatory receptors such as Chemotaxis Inhibitory Protein of *S. aureus* (CHIPS) (9), Formyl Peptide receptor-like 1 inhibitory protein (FLIPr), and the FLIPr homologue FLIPr-like (FLIPrL)(10, 11). Other immune evasion proteins serve as toxins that use surface receptors to specifically lyse leukocytes, such as the bi-component toxins (PVL, LukAB, LukED) (12–14) and phenol soluble modulins (15). Another group of secreted proteins, of which many are involved in immune evasion, are the Staphylococcal superantigen-like proteins (SSLs) (16).

SSLs are a family of 14 proteins with structural similarity to Staphylococcal superantigens but lack the functional T-cell receptor binding domain and therefore exhibit no superantigenic activity (17). Moreover, structurally, the C-terminal **β**-grasp domain of these SSL proteins show homology to other staphylococcal immune evasion proteins like CHIPS. SSL1 to SSL11 are encoded on staphylococcal pathogenicity island 2 whereas SSL12, SSL13 and SSL14 are found on the immune evasion cluster 2 (IEC-2) (4, 18). The SSL gene cluster is conserved in all human and animal isolates of *S. aureus* examined to date, indicating that it is very stable and evolutionary important cluster for the organism (18–20). Furthermore, antibodies against the SSLs are detected in human serum, indicating that they are expressed *in vivo* and may play a role during infection (20, 21). Even though the SSLs are highly conserved and involved in innate immune evasion, they have distinct functions (17). It was reported previously that several SSL members located on the main cluster (SSL3, SSL5, SSL6, SSL7 and SSL10) are involved in inhibition of host immune responses (22–25). SSL3 and SSL4 have been described as Toll-like receptor 2 (TLR2) inhibitors and prevent neutrophil activation (26, 27). SSL5 interacts with neutrophil surface receptor CD162 and reduces neutrophil migration (7, 23). SSL6 was identified to interact with CD47 by screening a *S. aureus* secretome phage display library for binding to isolated human neutrophils (2). SSL7 binds to complement C5 and therefore prevents C5a production (28). In addition, SSL7 and SSL10 are associated with blocking complement activation by targeting IgA and IgG respectively (28, 29). In contrast, none of the SSLs on the minor cluster (SSL12-14) have been functionally characterized.

In this study, we set out to identify new *S. aureus* proteins that interact with human neutrophils using a *S. aureus* secretome phage display library. In combination with Whole Genome Sequencing (WGS), SSL13 was identified to bind human neutrophils. We show that binding to human neutrophils is formyl-peptide receptor 2 (FPR2) dependent. Through this interaction, SSL13 activates neutrophils and acts as a chemoattractant. Furthermore, SSL13 activated neutrophils exhibit induced oxidative burst and degranulation. In contrast to many other immune evasion proteins that inhibit immune responses, we identified SSL13 as a chemoattractant and a neutrophil activator that acts via the FPR2.

## Experimental procedures

### Ethics statement

Informed consent was obtained from all subjects in accordance with the Declaration of Helsinki. Approval was obtained from the medical ethics committee of the University Medical Center Utrecht ((METC-protocol 07-125/C approved March 01, 2010; Utrecht, The Netherlands). The use of animals was approved by the National Ethical Committee for Animal Experiments (permit no AVD115002016565) and conducted according to local regulations.

### Reagents and antibodies

Monoclonal antibody (mAb) anti-His Tag (clone AD1.1.10, FITC-labeled) was purchased from LS Biosciences, and anti-CD62L (clone Dreg-56, FITC-labeled) and anti-CD11b (clone ICRF44, APC-labeled) were purchased from BD. The peptide MMK-1 (H-LESIFRSLLFRVM-OH) was synthesized by Sigma, and WKYMVM was synthesized by Bachem AG(Switzerland). WRWWWW-NH2 (WRW4) and Pertussis toxin were purchased from Tocris. Formyl-methionyl-leucyl phenylalanine (fMLP), TNF-α and cytochalasin B were from Sigma-Aldrich. Fluo-3-AM (acetoxymethyl ester) and Calcein-AM were purchased from Thermo Fisher.

### Cloning, expression, and purification of recombinant proteins

FLIPr, FLIPr-Like and N-terminal His-tag labeled SSL13 (His-SSL13) were cloned, expressed and purified as described (11, 30). For SSL13, primers were designed without signal peptide according to the published sequence of the gene NWMN_1076 for cloning into modified N-His6-TEV-(g)-pRSET vector (30). SSL13 was amplified from genomic DNA of *S. aureus* subsp. *aureus* strain Newman using the following primers: 5’-CG<u>GGATCC</u>CAATTTCCTAATACACCTATC-3’ and 5’-ATAT<u>GCGGCCGC</u>TTAGTTTGATTTTTCGAG-3’. Restriction enzyme recognition sites are underlined. Recombinant protein was generated in *E.coli* Rosetta Gami(DE3) plysS by induction with 1 mM Isopropyl β-D-1-thiogalactopyranoside (Roche). His-tagged protein was isolated under native purification conditions using a 5ml HiTrap chelating HP column (GE Healthcare) with an imidazole gradient (10-250 mM; Sigma-Aldrich). The purified protein was analyzed on a 12.5% SDS-PAGE gel and showed one band corresponding to a mass of 26.8 kD (Fig. S1). For direct fluorescent labeling, His-SSL13 was mixed with 0.1 mg/ml FITC (Sigma-Aldrich) in 0.1 M carbonate buffer (pH 9.5) for 1 h at 4°C and subsequently separated from free FITC by overnight dialysis against PBS.

### Cells

Human leukocytes were isolated from human heparinized blood as described (2) and suspended in RPMI-1640 supplemented with 20 mM Hepes (Gibco) containing 0.05% HSA (Sanquin). HL-60 cells were purchased from ATCC, HL-60 cells stable transfected with the human-FPR2 (HL-60/FPR2), were kindly provided by F. Boulay (Laboratoire Biochimie et Biophysique des Systemes Integres, Grenoble, France). Cells were cultured in RPMI-1640 supplemented with 10% fetal bovine serum (FCS), 100 μg/ml streptomycin, 100 units/ml penicillin.

### Phage library construction and phage production

A *S. aureus* secretome phage display library was created as described earlier (2). Briefly, genomic DNA from *S. aureus* strain Newman was mechanically fragmented and fragments were cloned into the pDJ01 secretome phagemid vector (2) and transformed into TG1 *E. coli*. Phages lacking an active pIII protein were produced overnight by co-infection with Hyperphage^®^ helper phages (Progen) at a multiplicity of infection of 10. Phages were purified and concentrated using PEG precipitation and resuspended in PBS to yield a final concentration of 2 x 10^11^ phages/ml.

### Phage selection on isolated human neutrophils

1 ml of phage library was mixed with 1 ml isolated human neutrophils (1 x 10^7^ in RPMI1640-0.05%HSA) and incubated on ice with gentle shaking for 30 min. Cells were washed twice by adding 50 ml cold RPMI-HSA and spinning down. Phages were eluted using 500 μl glycine 0.05M, pH2 for 5 min after which 62.5 μl neutralization buffer (2 M Tris-HCL pH 8.4) was added. Cells and cell debris were removed by centrifugation and phages were precipitated using 200 μl of 20% PEG/2.5 M NaCl for 30 min at room temperature. Sample was centrifuged at 14.000 rpm in an eppendorf centrifuge for 10 min at 4°C and supernatant was discarded. The pellet was suspended in 100 μl iodide buffer (10mM Tris-HCL, 1mM EDTA, 4M NaI, pH8) to disrupt the phage coat proteins and release the DNA. DNA was precipitated by adding 250 μl of 100% ethanol and incubated for 30 min at room temperature. Sample was centrifuged at 14.000 rpm in an eppendorf centrifuge for 10 min at 4°C after which the supernatant was discarded and the pellet containing the single stranded phage DNA was washed with 70% ice cold ethanol and dried to the air. The non-selected phage library was taken as a control.

### Phage library sequencing

Since the phage library was created using a pIII deficient helper phage, it consists of non-infectious phage particles. Therefore traditional phage selection with multiple rounds of selection and amplification is not possible and the library was analyzed by genome sequencing using the Illumina MiSeq System. In order to add the MiSeq adapters to the isolated phage DNA, a PCR reaction was performed on the precipitated DNA using Phusion^®^ HF polymerase (New England Biolabs), according to the manufacturer’s recommendations. The primers were designed for compatibility with the Illumina MiSeq v2 sequencing kit. (Table S1 for primer sequences). The PCR product was purified using gel purification on an Ultra-pure 2% agarose gel and the purified DNA was quantified on a Qubit 4 fluorometer (Thermo Fischer Scientific). The purified sample was run on a 1% agarose gel to determine purity and determine mean fragment size.

Sequencing was performed by loading 3pM of the library onto a MiSeq v2 2x250bp sequencing kit and ran on an Illumina MiSeq System according to manufacturer’s instructions. Sequence data was deposited in ENA under study accession number: PRJEB26168.

### His-SSL13 binding assay

To determine the binding of His-SSL13 to human leukocytes, a mixture of isolated neutrophils and mononuclear cells at 5 x 10^6^ cells/ml was incubated with increasing concentrations of His-SSL13 for 30 min at 4°C while gently shaking. Cells were washed and incubated with FITC-labeled anti-His tag mAb while shaking. Cells were washed and resuspended in buffer containing 1% paraformaldehyde (PFA). The fluorescence was measured on a FACSVerse flow cytometer, and the different leukocyte populations (neutrophils, monocytes and lymphocytes) were identified based on forward and sideward scatter parameters.

To determine the binding of His-SSL13 to HL-60 cells, 5x10^6^ cells/ml HL-60 cells were incubated with FITC-labeled SSL13 (SSL13-FITC) for 30 min at 4°C while shaking. Cells were washed and resuspended in buffer with 1% PFA. The fluorescence was measured by flow cytometry, and cell populations were identified based on forward and sideward scatter parameters excluding debris and death cells.

### CD11b and CD62L expression on neutrophils

Neutrophils (5 x 10^6^ cells/ml) were incubated with different concentrations SSL13 for 30 min at 37°C. Subsequently, the cells were put on ice and incubated with anti-CD11b and anti-CD62L mAb for 45 min on ice. Cells were washed and fixed with 1% PFA in buffer. Expression of CD11b and CD62L was measured on a flow cytometer and data expressed relative to the buffer treated cells.

### Calcium flux in neutrophils and HL-60 cells

Calcium flux with isolated human neutrophils and HL-60 cells was performed in a flow cytometer as previously described (31). Briefly, cells at 5 x 10^6^ cells/ml were labeled with 0.5 μM Fluo-3-AM ester, washed and resuspended to a concentration of 1x10^6^ cells/ml. To measure cells continuously and be able to add stimulus without interruption in the FACSVerse flow cytometer, the Eppendorf tube adapter was used without tube while sampling cells from a 96-well plate on an elevated platform. Stimuli were added in a 1/10^th^ sample volume after a 10 seconds baseline recording and calcium flux monitored for 50 seconds post stimulation. Samples were analyzed after gating neutrophils, thereby excluding cell debris and background noises. Calcium flux was expressed as difference between baseline fluorescence (mean of time point 3 till 8 sec) and after addition of stimulus (mean of time point 20 till 60 sec).

### Chemotaxis

Neutrophil migration was measured in a 96-multiwell transmembrane system (ChemoTX; Neuro Probe) using an 8 μm pore size polycarbonate membrane (32). Cells were labeled with 2 μM calcein-AM for 20 min, and resuspended to a concentration of 2.5x10^6^ cells/ml in HBSS with 1% HSA. Wells were filled with 29 μl of chemoattractant, and the membrane holder was carefully assembled. Cells were pre-incubated with or without FLIPr and 25 μl was placed as a droplet on the membrane. After incubation for 30 min at 37°C in a humidified 5% CO_2_ atmosphere, the membrane was washed extensively with PBS to wash away the non-migrating cells, and the fluorescence was measured in a fluorescence plate reader (CLARIOstar; BMG LABTECH) using 483 nm excitation and 530 emission filters. Percentage migration was calculated relative to wells containing the maximum number of 25 μl cells.

### Myeloperoxidase (MPO) release

Neutrophils were treated for 10 min with cytochalasin-B and TNF-alpha with gently shacking, and without wash, subsequently incubated with buffer only, SSL13 or fMLP. Cells were centrifugated at 500 x g for 10 min and supernatant collected for MPO activity measurement (33). Therefore, 10 μl sample was mixed with O-Dianisidine substrate and H_2_O_2_ in phosphate buffer at pH 6.0 and measured for 30 min at 37°C in a plate reader (FLUO star Omega) at 450 nm.

### Neutrophil oxidative burst assay

Horseradish peroxidase (HRP) and Isoluminol were used as a sensitive measure of the human neutrophil oxidative burst as described (34, 35). In white 96-well microtiter plates, a 150 μl reaction mixture of 6.25 x 10^4^ neutrophils per well in IMDM buffer with 0.1% HSA plus 50 μM Isoluminol and 4 U/ml HRP was equilibrated for 5 min. Subsequently concentrated stimulus was added to activate the NADPH-oxidase and emitted light immediately recorded continuously for 15 min in a Luminometer (Berthold) at 37°C. Data are expressed as relative light units (RLU).

### Mouse Experiments

In the mouse peritonitis model, 100 μg protein in 0.5 ml PBS was injected into the peritoneum of 6-to 8-week-old female CD-1 mice. At 4 hours later, the mice were euthanized by cervical dislocation and abdominal cavities washed with two times 5 ml of RPMI medium containing 0.1% HSA and 5mM EDTA. In total 8 to 9 ml of peritoneal fluid was recovered and centrifuged at 1200 rpm for 10 min to collect the exudate cells. Cell pellets were resuspended in 500 μl buffer and counted with trypan blue in a TC20 automated cell counter (BioRad). Before immuno staining, cells were first preincubated with 100 μg/ml normal goat IgG for 15 min. We stained the samples with APC-conjugated antibody to mouse CD45 (leukocytes marker), PE-conjugated antibody to mouse Gr1 (neutrophil marker), and FITC-conjugated antibody to mouse F4/80(macrophage marker). Samples were analyzed on a flow cytometer.

Mouse neutrophils were isolated from bone marrow as described previously (36). Briefly, a bone marrow cell suspension was collected by flushing the femurs and tibias with 10 ml of cold HBSS + 15 mM EDTA + 30 mM Hepes + 0.1% HSA. A two-layer Percoll density gradient (2 ml each in PBS) composed of 81% and 62.5% was used to enrich neutrophils from the total leucocyte population. Interphase between between 62.5% and 81% was collected. Cells were washed once with buffer and resuspended in PRMI1640 with 0.1% HSA.

Calcium fluxes in mouse neutrophils were determined as described above for human neutrophils with final concentrations of 10, 3, 1, 0.3, 0.1 and 0.03 nM of WKYMVM and 1000, 300, 100, 30, 10 and 3 nM of SSL13.

Mouse neutrophil binding assays were conducted essentially as described for human neutrophils.

## Results

### Phage library sequencing and identification of immune evasion

The sequencing run produced a total of 1,396 and 23,411 paired-end reads for the unselected and selection library, respectively. These reads were then quality-trimmed using nesoni clip v. 0.128 with the following parameters: –adaptor-clip yes –match 10 –max-errors 1 –clip-ambiguous yes –quality 10 –length 150 (http://www.vicbioinformatics.com/software.nesoni.shtm). About 90% of the read pairs were retained and used for further analyses.

Quality-trimmed sequence reads were aligned to the Genbank database (accessed on July 20^th^, 2015) using BLAST+ 2.2.31. 3 sequences in the non-selected and 4 sequences in the selected library did not align with a *S. aureus* genome and were omitted from analysis. The read frequency was defined as the total count of identical reads. The total amount of unique sequences per annotated gene was defined as number of clones. The highest hit in the unselected library is annotated as a dUTPase with a read frequency of 14 all belonging to a single clone. The 96 reads with the highest read frequency after selection encode for 61 different proteins that are listed in table S2. There is a large increase in read frequency after selection. The highest read frequency with 883 reads encoding 7 unique sequences is annotated as a transmembrane protein involved in mannitol transport. The selection of transmembrane proteins when performing phage display selection on cells was also observed in earlier phage selections in our lab (data not shown). The presence of membranes appear to select for transmembrane domains especially transporter proteins like ABC-transporters. The second highest hit with 196 reads and 4 different clones identified the recently described *S. aureus* protein (SPIN) that binds neutrophil myeloperoxidase and promotes the intracellular survival of *S. aureus* after phagocytosis (30). Of the total of 61 identified proteins, 12 (20%) were described to play a role in host microbe interaction. Of these 11 were already functionally characterized and for 1 protein, SSL13, no known function has been described. The fact that SSL13 was identified in this selection suggests that it is involved in binding to neutrophils or its components.

### SSL13 specifically interacts with human neutrophils

To confirm that SSL13 interacts directly with human neutrophils, a three-fold dilution series of recombinant SSL13 with an N-terminal His tag was incubated with human leukocytes isolated from healthy donors. His-tagged SSL7 and SSL5 were included as negative and positive control neutrophil-binding proteins respectively (7, 37). We observed that SSL13 interacts with human neutrophils and monocytes in a dose dependent manner, but no significant binding was observed to lymphocytes (Fig. 1A-C).

**Fig 1.**
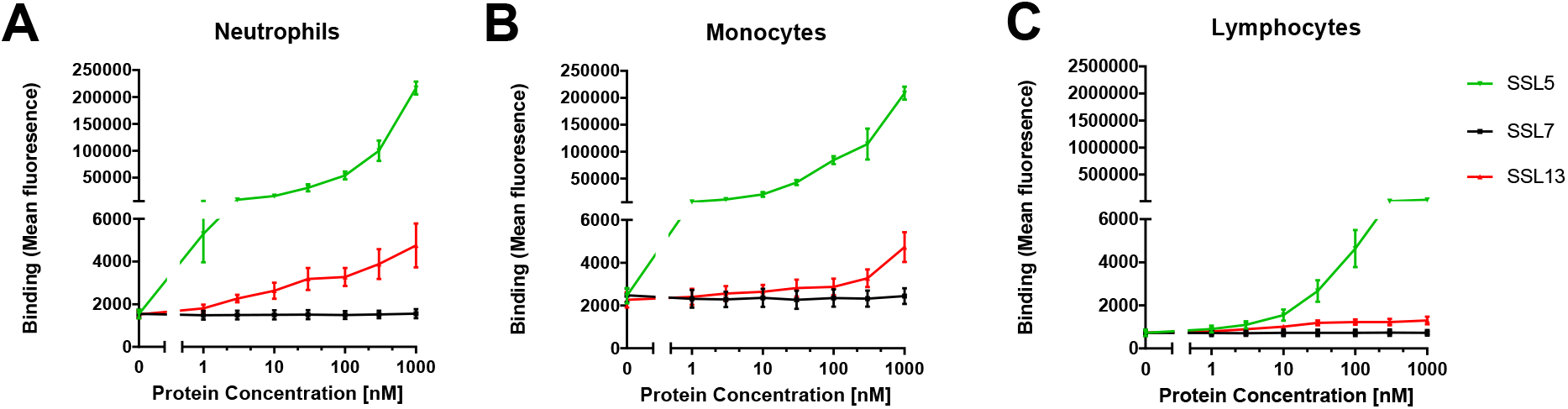
SSL13 binds to human neutrophils, monocytes, but not lymphocytes. Peripheral blood leukocytes were incubated with a three-fold dilution series of His-SSL13 for 30 min at 4 °C. Binding was detected with anti-His-FITC and analyzed by flow cytometry. The different cell populations were identified based on scatter parameters. His-SSL5 and His-SSL7 are positive and negative controls for binding respectively (A, B and C). Error bars are SEM of three biological replicates analyzed in duplicate.

Interestingly, binding experiments conducted at 37°C indicated that SSL13 activates neutrophils as shown by an increase in forward scatter compared with untreated cells (no protein) (38, 39) (Fig. S2 A-C). Activation of neutrophils generally alters the surface expression of major cell adhesion molecules, e.g. up-regulation of CD11b and down-regulation of CD62L (4). The effect of SSL13 on CD11b and CD62L expression was evaluated by flow cytometry. We observed that SSL13 enhanced the surface expression of CD11b and simultaneously down-regulated the expression of CD62L in a dose-dependent manner (Fig. 2A-B). In addition to the altered expression of surface adhesion molecules, activated neutrophils also exhibit intracellular release of calcium (40). We therefore measured the intracellular release of calcium after neutrophil exposure to a range of SSL13 concentrations (23-740 nM). In concordance with the cell receptor expression assay, our calcium flux data showed that SSL13 induces a transient dose-dependent release of Ca^2^+ in neutrophils (Fig. 2C-D). Degradation of SSL13 by proteinase K completely abolished the neutrophil activation indicating that the observed activation is not caused by a non-protein contaminant in the SSL13 preparation (Fig. S3 A-D). To conclude, SSL13 specifically binds, and activates human neutrophils.

**Fig 2.**
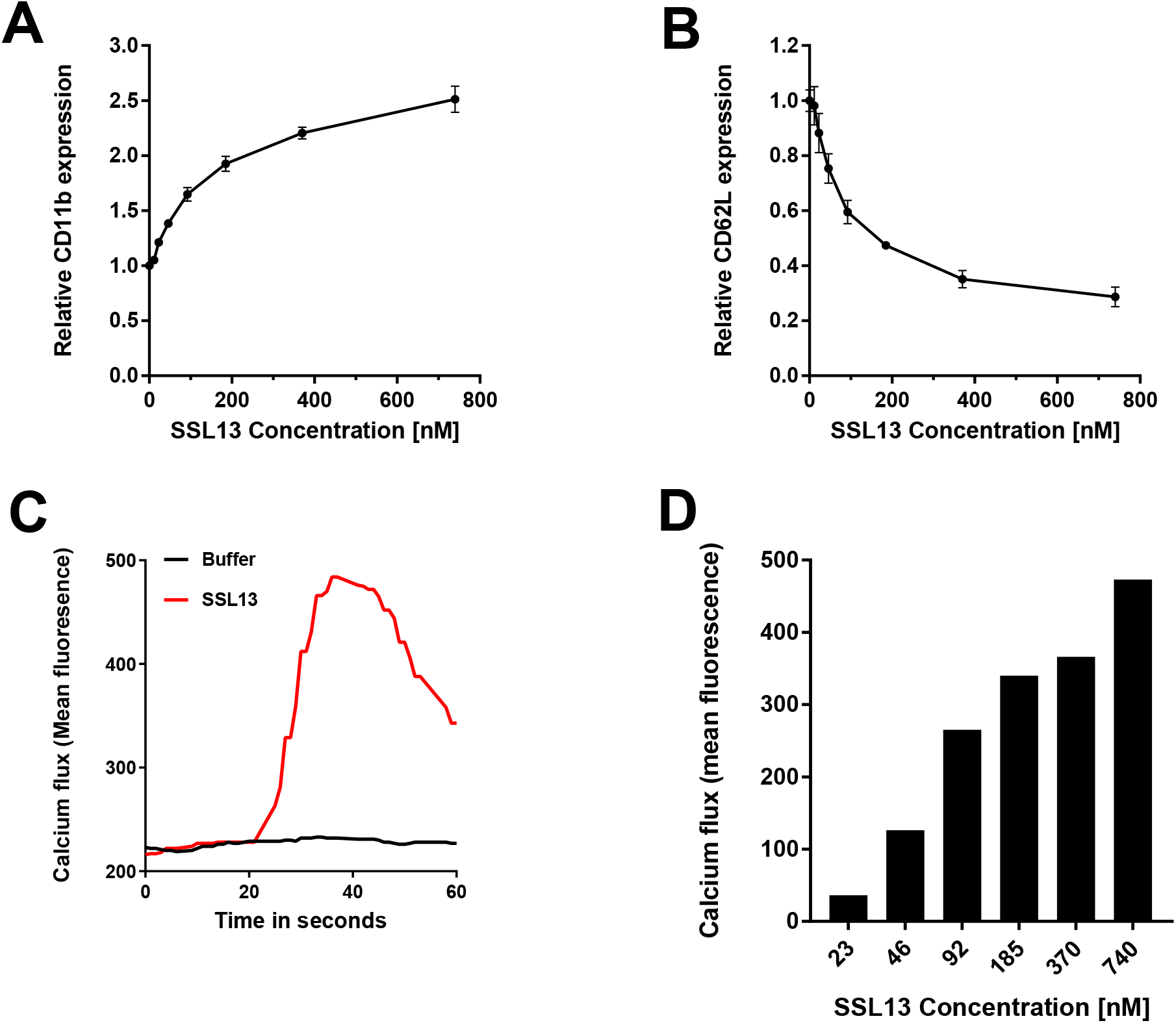
SSL 13 activates human neutrophils. (A and B) Activation of isolated human neutrophils by increasing concentration of SSL13. Increased CD11b expression (A) and decreased CD62L expression (B) are markers for neutrophil activation. Data are mean fluorescence ± SEM of three independent experiments. (C and D) Addition of SSL13 induces cell activation measured as a transient release of intracellular calcium (C). This effect is concentration dependent (D). Data are from one representative experiment.

### SSL13 specifically binds and activates formyl peptide receptor 2

As SSL13 induced a rapid and transient release of intracellular Ca^2+^, we examined whether SSL13 acts through a G protein-coupled receptor (GPCR) (41). Pertussis toxin (PTX) is a general antagonist of GPCR activation, and therefore blocks the release of intracellular Ca^2+^ (42). For this purpose, neutrophils were preincubated with or without PTX for 1 h at 37°C with CO_2_, and then stimulated with 370 nM SSL13 or fMLP as a reference PTX-sensitive stimulus (43, 44). Fig. 3A shows that PTX can block both SSL13 and fMLP induced neutrophil activation, which confirms that SSL13 utilizes a PTX-sensitive GPCR to induce this response.

**Fig 3.**
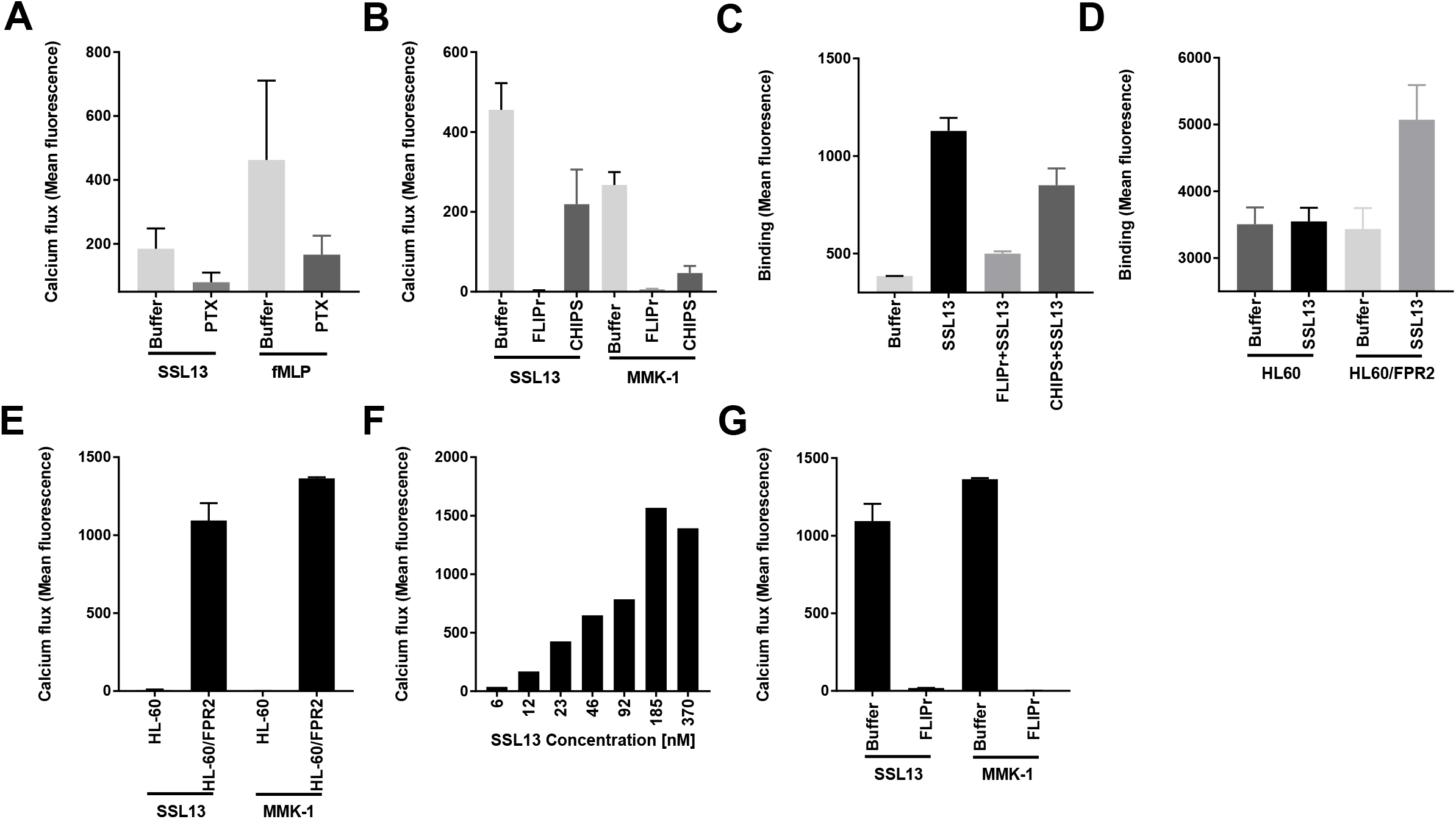
SSL13 specifically binds and activates human Formyl Peptide Receptor 2. (A) Neutrophils were preincubated with or without 3μg/ml PTX for 60min at 37 °C with CO2 and then labeled with Fluo-3 AM. Neutrophil stimulation by SSL13 is sensitive to PTX. fMLP is the control ligand of FPR1, which is also sensitive to PTX. (B) Human neutrophil stimulation by SSL13 is inhibited by FLIPr, not CHIPS. (C) SSL13 specific binding to human neutrophils is blocked by FLIPr, not CHIPS. Data represent means ± SEM of three independent experiments. (D) SSL13 specifically binds to FPR2 transfected HL60 cells(HL60/FPR2), but not to control HL60 cells. (E) SSL13 induces profound calcium fluxes in HL60/FPR2 cells, but not in empty HL60 cells. MMK-1 is a synthetic control ligand of FPR2. (F) HL60/FPR2 cells stimulation by SSL13 is concentration dependent. (G) HL60/FPR2 cells stimulation by SSL13 is sensitive to FPR2-specific inhibitor FLIPr. Data are mean fluorescence ± SEM of three experiments.

To further investigate the responsible receptor, a set of well-characterized agonists and antagonists of neutrophil GPCRs were tested, including those for formyl peptide receptor 1 (FPR1) and 2 (FPR2), Leukotriene B4 receptor, platelet activating factor (PAF) receptor, Complement C5a receptor, and the IL-8 receptors CXCR1 and CXCR2. We found that the FPR2 antagonists FLIPr inhibited the SSL13 induced calcium mobilization as well as the binding to human neutrophils (Fig. 3B-C). Although FLIPr also slightly inhibits the FPR1 activation, the control protein CHIPS, that specifically inhibits FPR1 (11), had no effect on SSL13-mediated neutrophil activation (Fig. 3C). Together, these experiments indicate that SSL13 elicits calcium fluxes in human neutrophils via FPR2.

To further confirm that FPR2 is the receptor for SSL13, we used HL60 cells stably transfected with or without human FPR2 (45, 46). Binding of FITC-labeled SSL13 was only observed with HL60/FPR2 and not with control HL60 cells (Fig. 3D). Furthermore, in order to evaluate the role of FPR2 in recognizing SSL13, we analyzed the intracellular Ca^2+^ response to SSL13 of HL60 with or without FPR2. Fig. 3E shows that SSL13 induces a profound calcium flux in HL60/FPR2, but not in untransfected HL60 cells. The activation potential of SSL13 is comparable to the specific FPR2 agonistic peptide MMK-1 (Fig. 3E). Moreover, SSL13 activates the FPR2 transfected HL60 cells in a dose dependent manner (Fig. 3F). Finally, the induced calcium flux of the FPR2 transfected HL60 cells by SSL13 and MMK-1 can be inhibited by the FPR2-specific inhibitor FLIPr (Fig. 3G). These findings confirm that SSL13 specifically binds and activates cells via FPR2.

### SSL13 is involved in chemoattractant induced oxidative burst and degranulation of neutrophils

Triggering FPR2 induces many neutrophil effector functions, including chemotaxis, exocytosis and superoxide generation (47). To investigate whether SSL13 is a chemoattractant, neutrophil migration was measured in a 96-multiwell transmembrane system. Indeed, SSL13 stimulates chemotaxis of human neutrophils in a dose-dependent manner (Fig. 4A). Moreover, the SSL13 induced chemotaxis in human neutrophils can be blocked by the FPR2 antagonist FLIPr (Fig. 4B).

**Fig 4.**
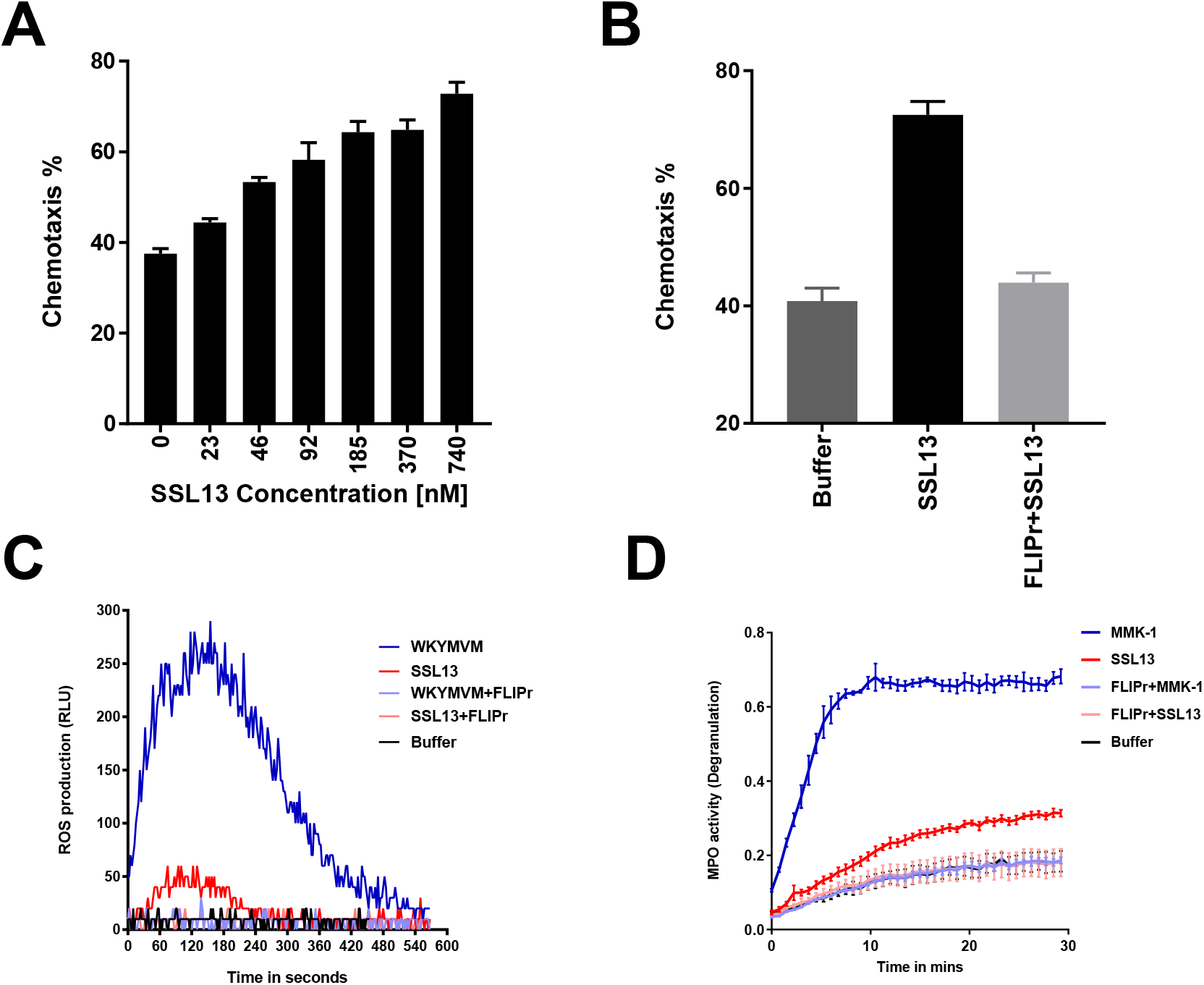
SSL13 is involved in chemoattractant induced oxidative burst and degranulation of neutrophils. (A) SSL13 stimulates chemotaxis in human neutrophils in a dose dependent manner. (B) SSLl3-induced chemotaxis of human neutrophils is inhibited by the FPR2 antagonist FLIPr. (C) SSL13 stimulates FPR2 induced oxidative burst. WKYMVM is a synthetic control ligand of FPR2. (D) SSL13 modestly induces neutrophil degranulation via FPR2. MMK-1 is a positive control. (A and B) data represent means ± SEM of three experiments. (C and D) data from a representative experiment.

A common feature of most GPCRs is that they not only strongly activate the chemotactic migration of neutrophils, but also trigger neutrophil oxidative burst and degranulation. To examine whether SSL13 is involved in FPR2-induced oxidative burst, a Reactive Oxygen Species (ROS) assay was performed. The peptides WKYMVM and MMK-1 can both induce FPR2-mediated ROS production, although WKYMVM is more potent and was therefore used as control in our experiment (48). Our data shows that SSL13 induced a modest oxidative burst compared with the control FPR2 specific peptide WKYMVM (Fig. 4C), but both SSL13- and WKYMVM-induced oxidative burst in human neutrophils could be blocked by FLIPr (Fig. 4C). Furthermore, we tested whether SSL13 could induce neutrophil degranulation by measuring myeloperoxidase (MPO) activity in stimulated cell supernatant. MPO is one of the most abundantly granule proteins in neutrophils and efficiently released into the extracellular space during degranulation (49). Indeed, SSL13 induces neutrophil degranulation (Fig. 4D). Taken together, the functional outcomes of SSL13-induced neutrophil activation include chemotaxis, ROS production and neutrophil degranulation pointing toward a pro-inflammatory response of neutrophils to this staphylococcal protein.

To test whether SSL13 could act intracellular and are produced by *S.aureus* after uptake by human neutrophils, we generated a GFP promoter construct. Since SSL13 is part of operon together with SSL12 and SSL14, the SSL12-13-14 promoter was cloned in front of GFP and transformed into *S.aureus* Newman. We did not observe expression of GFP under various standard culture conditions or after uptake of bacteria by phagocytes as seen with some other staphylococcal immune evasion proteins (SPIN) (data not show here) (33).

### SSL13 is not able to efficiently activate mouse neutrophils

Many other Staphylococcal immune evasion proteins show a high level of human specificity. In order to check the host-dependent activation of SSL13, we tested binding and activation of neutrophils isolated from mice bone marrow. SSL13 can induce activation of murine neutrophils as shown by calcium mobilization. Treating the cells with WRW4, a known inhibitor of mice FPR2 (50), prevented the SSL13-induced calcium flux. This indicates that the neutrophil activation by SSL13 happened in a murine FPR2 dependent manner (Fig. 5A), although much higher concentrations are needed as compared to human neutrophil activation (Fig. 5B). In contrast, the specific FPR2 agonistic peptide WKYMVM showed similar activation ability to both human and murine neutrophils (Fig. 5C). However, we are unable to detect any SSL13 binding to murine neutrophils (data not shown).

**Fig 5.**
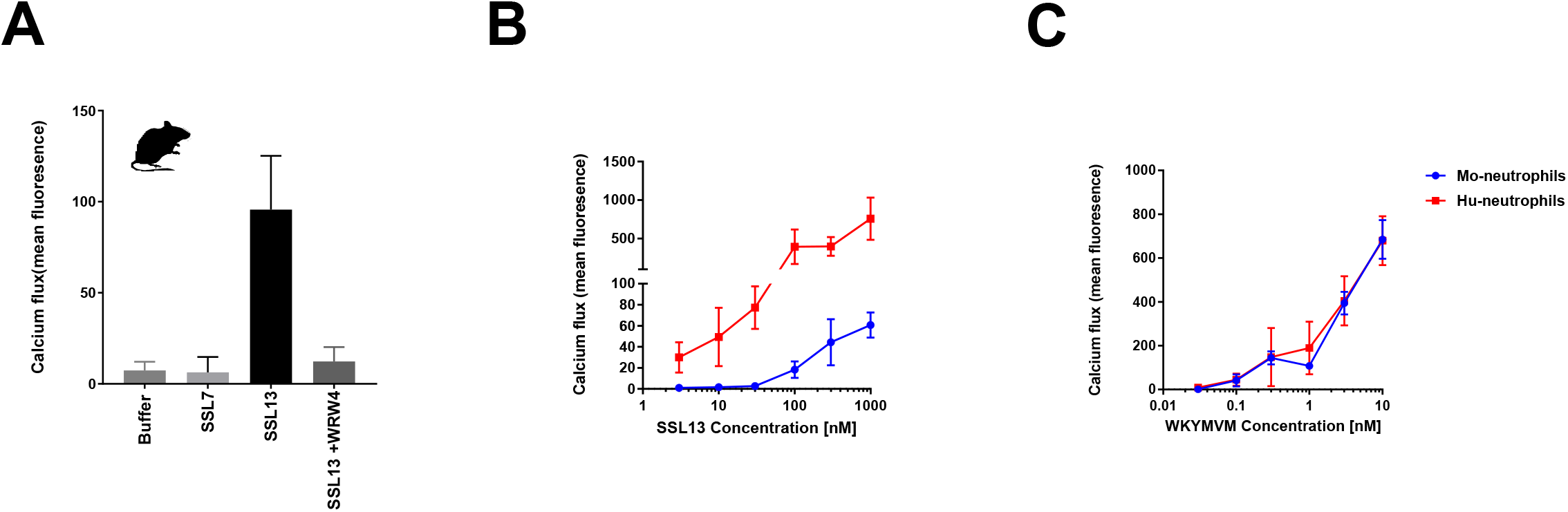
SSL13 is not able to efficiently activate mouse neutrophils. (A) SSL13 can induce activation of murine neutrophils, which can be inhibited by the mFPR2 antagonist WRW4. (B) SSL13-induced calcium fluxes in murine neutrophils are low compared to human neutrophils. (C) WKYMVM-induced calcium fluxes in murine neutrophils are similar to human neutrophils. Data are mean fluorescence ± SEM of three experiments.

Since there was a minimal but specific activation of mouse neutrophils, we tested whether SSL13 can provoke a neutrophil influx after injection of SSL13 into the mouse-abdominal cavity. We observed no increase in peritoneal neutrophil numbers at 4 h after intra-abdominal injection of 100 μg SSL13 (data not show). This indicates that SSL13 is highly adapted to specifically act on human neutrophils.

## Discussion

Previously, our group described a high-throughput binding selection strategy, phage display, to identify *S. aureus* immune evasion molecules. In this strategy, only secreted proteins of a bacterial genome are expressed on the surface of a filamentous phage, which is well suited to identify and characterize immune evasion proteins (2). Traditional phage selection strategies involve multiple rounds of selection and amplification and selecting single clones for sequencing and further analysis. Whole genome Illumina sequencing allows to analyze a phage library after only a single round of selection omitting library amplification that would undoubtedly lead to additional selection bias. Using this strategy we identified 12 proteins involved in host microbe interaction or immune evasion in a single round of selection indicating the enormous potential of this strategy. Furthermore, 8 conserved hypothetical proteins identified need further characterization, and may also be involved in host microbe interaction. The identification of SSL13, a protein with previously unknown function, in this phage selection suggested an interaction between SSL13 and neutrophils.

The SSLs are a family of 14 secreted proteins which were previously demonstrated to modulate immune evasion (3, 4, 17). Genetic analyses of 88 clinical *S. aureus* strains revealed that the genes encoding SSL12, SSL13, and SSL14 are conserved among all strains (51). We also confirm that SSL13 is produced *in vivo* as antibodies against those proteins can be detected in human serum (Fig. S4). Furthermore, in sharp contrast to the SSLs located on SPI-2 that all have their own promoter, SSL12-13-14 share a single promotor. Our hypothesis is that SSL12-13-14 may be produced simultaneously by *S. aureus* under certain conditions and that their function is linked. We show that SSL13 interacts with human neutrophils via FPR2. This interaction leads to activation and chemotaxis. We propose that the chemotactic property of SSL13, via FPR2, is important during early infection with *S. aureus* to lure neutrophils to the site of infection. Further studies should resolve the function of SSL12 and SSL14, which are simultaneously expressed under the same promotor. Just like the *S. aureus* bi-component toxin PVL requires LukS-PV and LukF-PV to properly lyse neutrophils (12), SSL12-13-14 might require the presence of all three proteins to elicit its maximum potential in immune modulation. Expression and secretion of SSLs under standard culture conditions is very limited and only low amounts of protein can be found in the cell culture supernatant. Previous research showed that there is an impressive upregulation and expression of some SSLs under different stress conditions (51). We indeed could not observe SSL13 expression by using GFP reporter construct in standard cell culture or after uptake by neutrophils.

SSL13 is not the only secreted molecule from *S. aureus* that is able to activate neutrophils. Phenol Soluble Modulins (PSMs), which are small peptides secreted by *S. aureus*, and have completely different structure compared to SSL13, are known to activate and attract both human and mice neutrophils via FPR2 (15, 50, 53). In addition to this, micromolar concentrations of PSM have cell lytic activity which is independent from FPR2. Serum can fully block PSMs functions in both the cell lysis and FPR2-mediated neutrophil activation (15). However, SSL13 activity was not inhibited by serum and SSL13 is not cytotoxic to neutrophils (Fig. S5). In contrast to PSMs, SSL13 showed a high degree of human specificity that was not able to efficiently activate mouse neutrophils.

FLIPr and its homologue FLIPr-like (FLIPrL) are located on the same IEC-2 cluster as SSL13, which are found in many, but not all, human *S. aureus* isolates (51). SSL13 is a neutrophil chemoattractant and activator that acts via the FPR2, whereas FLIPr and FLIPrL bind and inhibit FPR2 signaling function (10, 11). This may contribute to the ability of *S. aureus* to adjust a favorable balance between neutrophil activation and inhibition. Similar to other staphylococcal immune evasion proteins, many of the SSL proteins harbor several distinct functions. Therefore, it is not unlikely that SSL13 may has another unique function beyond activating FPR2 signaling. To conclude, SSL13 is a unique SSL member that does not belong to the immune evasion class, but is a pathogen alarming molecule acting on FPR2.

## ACKNOWLEDGMENTS

This work was supported by a grant from the program of China Scholarships Council (No. 201406170045). We thank Vincent P van Hensbergen for sharing expertise and reviewing the manuscript.

---

**Table S1.**
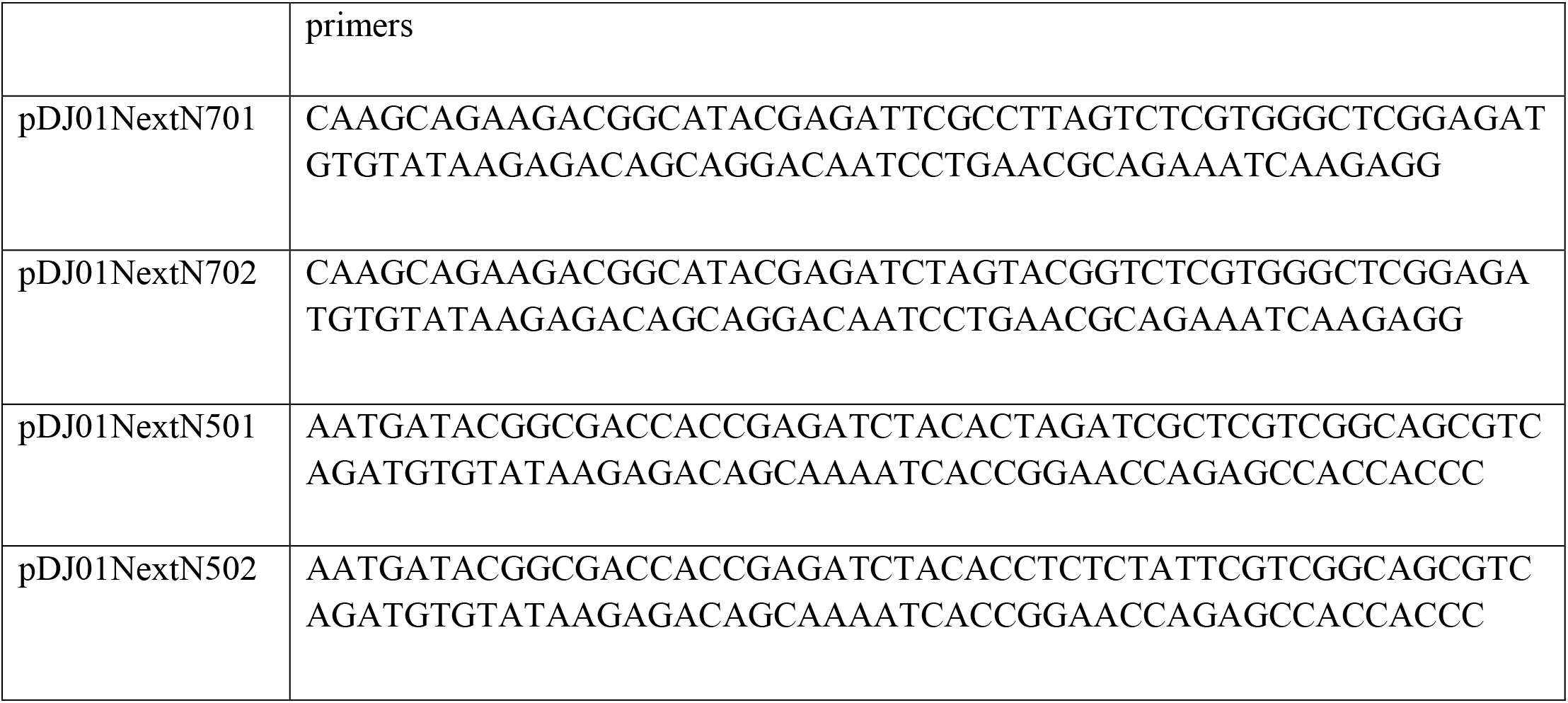
Primers for genome sequencing.

**Table S2.**
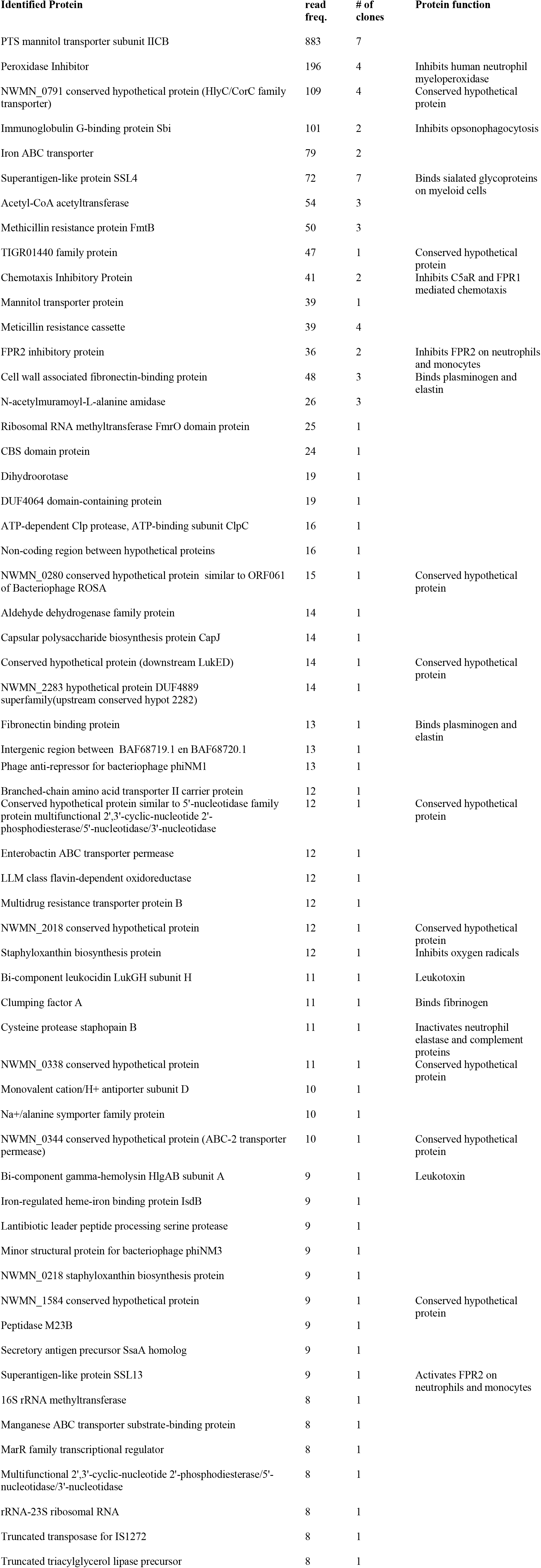
Proteins with the highest read frequency after phage display selection. A *Staphylococcus aureus* phage display library was selected for binding against isolated human neutrophils. The selected library was analyzed by whole genome sequencing. Table shows the top hits after selection based on read frequency and grouped by number of different clones. Function of previously characterized immune evasion proteins and hypothetical proteins are listed.

| Identified Protein | read<br>freq. | # of<br>clones | Protein function |
| --- | --- | --- | --- |
| PTS mannitol transporter subunit IICB | 883 | 7 |  |
| Peroxidase Inhibitor | 196 | 4 | Inhibits human neutrophil<br>myeloperoxidase |
| NWMN_0791 conserved hypothetical protein (HlyC/CorC family<br>transporter) | 109 | 4 | Conserved hypothetical<br>protein |
| Immunoglobulin G-binding protein Sbi | 101 | 2 | Inhibits opsonophagocytosis |
| Iron ABC transporter | 79 | 2 |  |
| Superantigen-like protein SSL4 | 72 | 7 | Binds sialated glycoproteins<br>on myeloid cells |
| Acetyl-CoA acetyltransferase | 54 | 3 |  |
| Methicillin resistance protein FmtB | 50 | 3 |  |
| TIGR01440 family protein | 47 | 1 | Conserved hypothetical<br>protein |
| Chemotaxis Inhibitory Protein | 41 | 2 | Inhibits C5aR and FPR1<br>mediated chemotaxis |
| Mannitol transporter protein | 39 | 1 |  |
| Meticillin resistance cassette | 39 | 4 |  |
| FPR2 inhibitory protein | 36 | 2 | Inhibits FPR2 on neutrophils<br>and monocytes |
| Cell wall associated fibronectin-binding protein | 48 | 3 | Binds plasminogen and<br>elastin |
| N-acetylmuramoyl-L-alanine amidase | 26 | 3 |  |
| Ribosomal RNA methyltransferase FmrO domain protein | 25 | 1 |  |
| CBS domain protein | 24 | 1 |  |
| Dihydroorotase | 19 | 1 |  |
| DUF4064 domain-containing protein | 19 | 1 |  |
| ATP-dependent Clp protease, ATP-binding subunit ClpC | 16 | 1 |  |
| Non-coding region between hypothetical proteins | 16 | 1 |  |
| NWMN_0280 conserved hypothetical protein similar to ORF061<br>of Bacteriophage ROSA | 15 | 1 | Conserved hypothetical<br>protein |
| Aldehyde dehydrogenase family protein | 14 | 1 |  |
| Capsular polysaccharide biosynthesis protein CapJ | 14 | 1 |  |
| Conserved hypothetical protein (downstream LukED) | 14 | 1 | Conserved hypothetical<br>protein |
| NWMN_2283 hypothetical protein DUF4889<br>superfamily(upstream conserved hypot 2282) | 14 | 1 |  |
| Fibronectin binding protein | 13 | 1 | Binds plasminogen and<br>elastin |
| Intergenic region between BAF68719.1 en BAF68720.1 | 13 | 1 |  |
| Phage anti-repressor for bacteriophage phiNM1 | 13 | 1 |  |
| Branched-chain amino acid transporter II carrier protein | 12 | 1 |  |
| Conserved hypothetical protein similar to 5'-nucleotidase family protein multifunctional 2',3'-cyclic-nucleotide 2'-phosphodiesterase/5'-nucleotidase/3'-nucleotidase | 12 | 1 | Conserved hypothetical protein |
| Enterobactin ABC transporter permease | 12 | 1 |  |
| LLM class flavin-dependent oxidoreductase | 12 | 1 |  |
| Multidrug resistance transporter protein B | 12 | 1 |  |
| NWMN_2018 conserved hypothetical protein | 12 | 1 | Conserved hypothetical protein |
| Staphyloxanthin biosynthesis protein | 12 | 1 | Inhibits oxygen radicals |
| Bi-component leukocidin LukGH subunit H | 11 | 1 | Leukotoxin |
| Clumping factor A | 11 | 1 | Binds fibrinogen |
| Cysteine protease staphopain B | 11 | 1 | Inactivates neutrophil elastase and complement proteins |
| NWMN_0338 conserved hypothetical protein | 11 | 1 | Conserved hypothetical protein |
| Monovalent cation/H<sup>+</sup> antiporter subunit D | 10 | 1 |  |
| Na<sup>+</sup>/alanine symporter family protein | 10 | 1 |  |
| NWMN_0344 conserved hypothetical protein (ABC-2 transporter permease) | 10 | 1 | Conserved hypothetical protein |
| Bi-component gamma-hemolysin HlgAB subunit A | 9 | 1 | Leukotoxin |
| Iron-regulated heme-iron binding protein IsdB | 9 | 1 |  |
| Lantibiotic leader peptide processing serine protease | 9 | 1 |  |
| Minor structural protein for bacteriophage phiNM3 | 9 | 1 |  |
| NWMN_0218 staphyloxanthin biosynthesis protein | 9 | 1 |  |
| NWMN_1584 conserved hypothetical protein | 9 | 1 | Conserved hypothetical protein |
| Peptidase M23B | 9 | 1 |  |
| Secretory antigen precursor SsaA homolog | 9 | 1 |  |
| Superantigen-like protein SSL13 | 9 | 1 | Activates FPR2 on neutrophils and monocytes |
| 16S rRNA methyltransferase | 8 | 1 |  |
| Manganese ABC transporter substrate-binding protein | 8 | 1 |  |
| MarR family transcriptional regulator | 8 | 1 |  |
| Multifunctional 2',3'-cyclic-nucleotide 2'-phosphodiesterase/5'-nucleotidase/3'-nucleotidase | 8 | 1 |  |
| rRNA-23S ribosomal RNA | 8 | 1 |  |
| Truncated transposase for IS1272 | 8 | 1 |  |
| Truncated triacylglycerol lipase precursor | 8 | 1 |  |

